# The Y-chromosome clarifies the evolutionary history of *Sus scrofa* by large-scale deep genome sequencing

**DOI:** 10.1101/126060

**Authors:** Huashui Ai, Jun Ren, Junwu Ma, Zhiyan Zhang, Wanbo Li, Bin Yang, Lusheng Huang

## Abstract

The genetics and evolution of sex chromosomes are largely distinct from autosomes and mitochondrial DNA (mtDNA). The Y chromosome offers unique genetic perspective on male-line inheritance. Here, we uncover novel evolutionary history of *Sus scrofa* based on 205 high-quality genomes from worldwide-distributed different wild boars and domestic pig breeds. We find that only two haplotypes exist in the distal and proximal blocks of at least 7.7 Mb on chromosome Y in pigs across European and Asian continents. And the times of most recent common ancestors (T_MRCA_) within both haplotypes, approximately 0.14 and 0.10 million years, are far smaller than their divergence time of around 1.07 million years. What’s more, the relationship between Sumatran and Eurasian continent *Sus scrofa* is much closer than that we knew before. And surprisingly, European pigs share the same haplotype with many Chinese pigs, which is not consistent with their deep splitting status on autosome and mtDNA. Further analyses show that the haplotype in Chinese pigs was likely introduced from European wild boars via ancient gene flow before pig domestication about 24k years ago. Low mutation rates and no recombination in the distal and proximal blocks on chromosome Y help us detect this male-driven ancient gene flow. Taken together, our results update the knowledge of pig demography and evolution, and might shed insight into the genetics and evolution studies on chromosome Y in other mammals.

## Introduction

The pig (*Sus scrofa*) provides human a main source of animal protein and serves as a biomedical model for human disease (Groenen et al. 2012). Since pigs are usually raised to yield meat and feed a majority of the world population, lots of studies have been conducted to reveal the genetic mechanisms for the complex economic traits in pigs, such as growth and fatness (Andersson et al. 1994), meat quality (Milan et al. 2000; Ma et al. 2014) and reproduction (Uimari et al. 2011). On the other hand, pigs have experienced a long period of breeding in yards or fields adjunct to human agricultural societies; they evolved similar eating pattern and dietary structure to human beings (Fang et al. 2012). And pigs share high resemblances with humans in terms of anatomy and physiology. It is generally believed that pigs can be treated as the ideal animal model for studying human microbial infectious diseases (Meurens et al. 2012), or as the promising candidate for development of tissue engineering techniques and xenotransplantation (Eventov-Friedman et al. 2006; Ekser et al. 2012). Therefore pig, as an important domestic animal, is very close related to human. Elucidating pig evolutionary history increases our cognition to pigs, provides insights to the development of two foundational functions of pigs, and helps us better understand the demographic history of humans.

Currently, pig demography and evolutionary history have been adequately revealed by lots of great works, which mainly included three aspects of zooarchaeological analyses, mtDNA or ancient mtDNA studies, and autosomal genomic researches. It is widely accepted that *Sus scrofa* emerged in Island South East Asia (ISEA) during the climatic fluctuations of the early Pliocene about 3 to 4 million years ago (Mya) and over the past one million year colonized almost the entire Eurasian continent (Frantz et al. 2013; Groenen 2016). On the northern parts of Sumatran Island, one of Southeast Asian islands, a wild boar population is also found, which split from Eurasian wild boar around 1.6 to 2.4 Mya (Frantz et al. 2013; Groenen 2016). The European and Asian wild boar populations diverged around 1 Mya, and North and South Chinese *Sus scrofa* populations separated from each other during the Ionian stage approximately 0.6 Mya (Groenen et al. 2012; Frantz et al. 2013). Pigs were domesticated at least at two locations (Anatonia and China) ∼10k years ago (Larson et al. 2005; Larson et al. 2007; Frantz et al. 2013), and gradually formed a variety of breeds in Europe and Asia (Kijas and Andersson 2001; Wang et al. 2011; Ottoni et al 2013). While during and after domestication of pigs, long-term gene flows or hybridization between wild boars and domestic pigs have been evidenced (Giuffra et al. 2000; Frantz et al. 2015). Nowadays Eurasia has the most rich pig resources, and about one third of worldwide pig breeds have adapted to divergent environment of China (Wang et al. 2011). Recently, a complex pattern of admixture and introgression between European and Asian domestic pigs, and African and American feral pigs has been well documented, but most of pigs colonizing the American, African and Australian continents originate from two highly distinct source populations of European and Asian local pigs (Ramirez et al. 2009; White 2011; Noce et al. 2015).

These above studies about pig demography and evolution are mainly focused on zooarchaeological evidences and genetic variations on autosomes and mtDNA. Few works were conducted from the perspectives of whole sex chromosomes, especially for the Y-chromosome, possibly due to lacking of enough genomic information. On the X-chromosome, we have previously found an interesting event of possible ancient interspecies introgression by sequencing 69 Chinese local pigs (Ai et al. 2015), which made an important complement to pig evolutionary history. With the development of sequencing technology, 13 de novo assembled pig genomes (Fang et al. 2012; Groenen et al. 2012; Li et al. 2013; Vamathevan et al. 2013; Li et al. 2016) were available for public access. But the version of Build 10.2 reference genome was still widely used due to comprehensive gene annotation and rich variants information. More recently, an improved assembly and gene annotation of the pig X Chromosome and a draft assembly of the pig Y Chromosome (VEGA62) were also generated by sequencing BAC and fosmid clones from Duroc animals and incorporating information from optical mapping and fiber-FISH (Skinner et al. 2016). These improved assemblies provide us make a profound survey to the evolutionary history of pigs from the perspectives of sex chromosomes, especially from the unique genetic perspective on male-line inheritance.

In the current work, we obtained high-quality whole-genome sequence data of 202 pigs from divergent populations and three outgroups including *Phacochoerus africanus, Sus verrucosus* and *Sus celebensis*. Based on variants of the Y-chromosome, we presented a deep investigation for male-line evolutionary history in pigs from the global, with assist of autosomal and mtDNA information.

## Results

We have previously sequenced 69 Chinese typical indigenous pigs to reveal the genetic basis for porcine local climate adaptation and possible ancient interspecies introgression on the X-chromosome (Ai et al. 2015). Here we increase 104 pigs for next-generation sequencing, download public data of 39 pigs and 3 outgroups, and combine our previous samples; we make a high-quality dataset of 205 individuals (**Fig. 1A, Supplemental Table S1**). All data was mapped into the *Sus scrofa* reference genome (build 10.2) using BWA (Li and Durbin 2009). A total of 48,745,075 SNPs were identified in the 205 genomes (**Supplemental Table S2**) using Platypus (Rimmer et al. 2014) under the criterion of minor allele frequencies (MAF) greater than 0.001 and call rates greater than 80%. All male data was remapped into the Y-chromosome (VEGA62) using Bowtie2 (Langmead and Salzberg 2012). On the new version of chromosome Y, a total of 49,103 SNPs were detected in the males using Platypus under the above criterion of MAF > 0.001 and call rates > 80%. These SNPs were used for subsequent demographic and evolutionary analyses.

**Figure 1.**
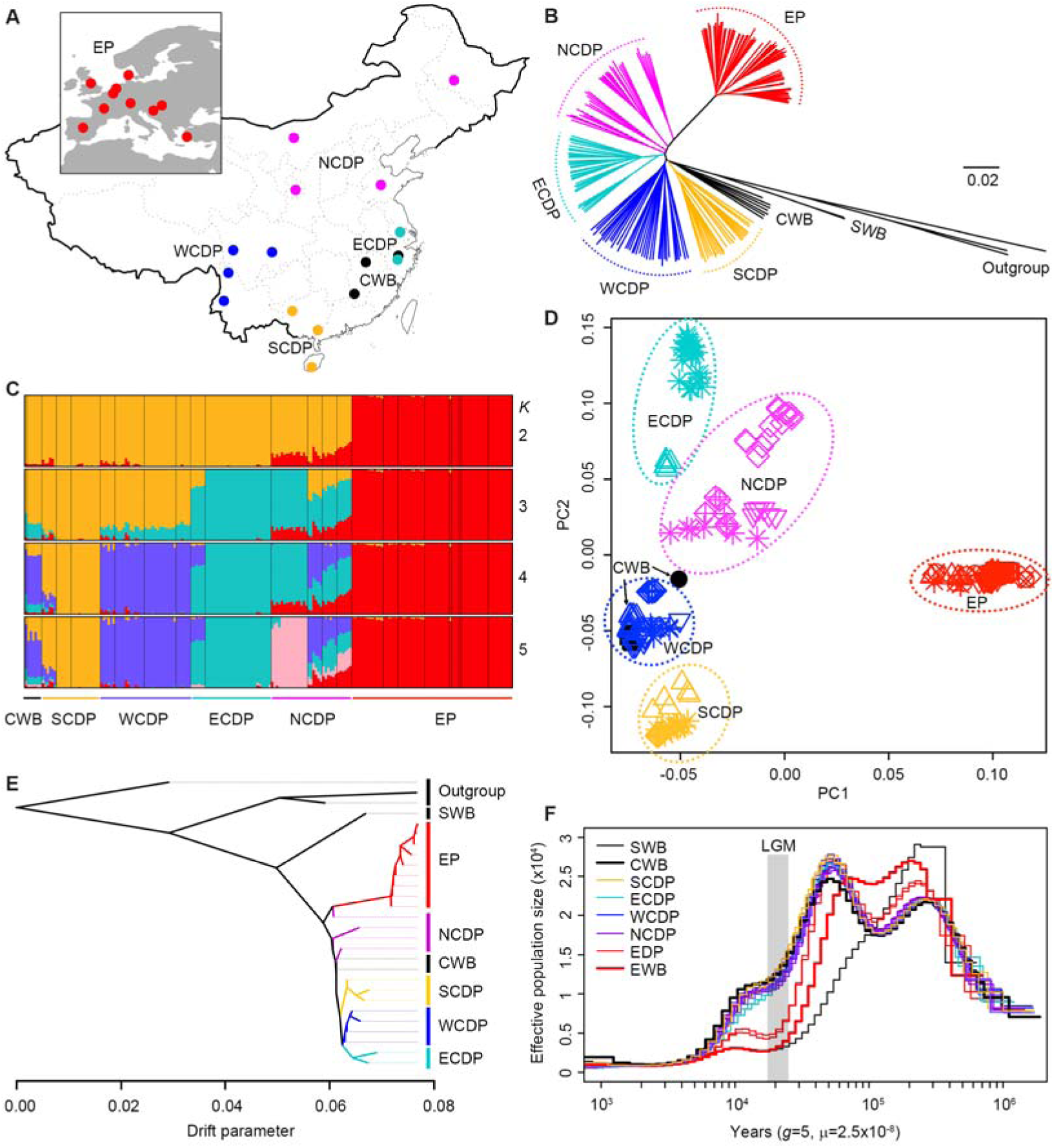
Demographic history of Eurasian pigs inferred using autosomal SNP data. (A) Geographical locations of Eurasian pigs analyzed in this study. EP, European pigs including Creole, Duroc, Iberian, Large White, Mangalica, Landrace, Pietrain and wild boars; ECDP, East Chinese domestic pigs including Erhualian and Jinhua; WCDP, West Chinese domestic pigs including Baoshan, Neijiang and Tibetan pigs in Yunnan and Sichuan provinces; SCDP, South Chinese domestic pigs including Bamaxiang, Luchuan and Wuzhishan; NCDP, North Chinese domestic pigs including Bamei, Hetao, Laiwu and Min; CWB, Chinese wild boars. (B) Neighbor joining phylogenic tree of all sequenced pigs. SWB, Sumatran wild boars. *S. celebensis* (Celebes wild boar), *S. verrucosus* (Java warty pig) and *P. africanus* (African warthlog) were used as outgroups. (C) ADMIXTURE analysis with *K* = 2-5. Colors in each column represent ancestry proportion. (D) Principal component analysis plots based on the first two principal components. (E) Relationships among Eurasian pigs inferred using Treemix. (F) Effective population sizes of Eurasian pigs inferred using MSMC. The period of the Last Glacial Maximum (LGM; ∼20,000 years ago) is shaded in grey.

### Demographic parameters inferred by autosomal data

We first constructed a neighbor-joining (NJ) tree for the above 205 animals using 47,009,938 SNPs on autosomes. All individuals from the same population gathered together, and European pigs defined a branch clearly separated from Chinese pigs in the NJ tree (**Fig. 1B**). Chinese wild boars clustered into one group, and Chinese domestic pigs were roughly categorized into four groups, corresponding to their geographical distributions: South China, West China, East China and North China (**Fig. 1B**). We then conducted principal components (Price et al. 2006), ADMIXTURE (Alexander et al. 2009) and Treemix (Pickrell and Pritchard 2012) analyses to assess population structure of these animals. European pigs formed an independent lineage that was genetically distinct from Chinese pigs in both the ADMIXTURE results (**Fig. 1C**) and the principal components plot (**Fig. 1D**). The Treemix analysis inferred the deep divergence not only between Sumatran pigs and Eurasian pigs, but also between European and Chinese pigs, and North Chinese pigs showed closer relationship with European pigs than other Chinese pigs (**Fig. 1E**). We further inferred the history of population size of Sumatran, Chinese and European pigs using the multiple sequentially Markovian coalescent method (Schiffels and Durbin 2014). Obviously, the demographic profile of Sumatran pigs largely differed from those of Chinese and European pigs. But from the ancient time of the Last Glacial Maximum to the recent 1000 years, the curve of Sumatran pigs almost overlapped with the one of European wild boars. Interestingly, the demographic profiles of Chinese pigs began to diverge from those of European pigs about 0.3 Mya (**Fig. 1F**), and had experienced less severe decline in population size during the Last Glacial Maximum, which is consistent with the previous reports (Groenen et al. 2012; Frantz et al. 2013). Notably, all the test pigs had experienced a bottleneck effect from ∼ 5000 to 2000 years ago, which was most severe than the past time. We hypothesized that the activities of human domestication or human hunting might contribute to the decease of pig or wild boar across the global.

Altogether, these results based on the autosomal data support the previous conclusion that Chinese and European pigs represent two genetically divergent ancestral populations and Sumatran wild boars are largely different with Eurasian pigs (Larson et al. 2005; Groenen et al. 2012; Frantz et al. 2013).

### Odd haplotype patterns of both sex chromosomes in Eurasian pigs

We then made a close examination on sex chromosomes using 1,730,532 and 49,103 SNPs on chromosomes X and Y, respectively. As expected, the low-recombinant low-mutation-rate region of 52 Mb (from 49 to 101 Mb on Build 10.2 reference genome) on the X chromosome exhibits three different major haplotypes: one found in European pigs and North Chinese wild boars; one found in pigs from South and West China; and the recombinant haplotype between the two haplotypes was observed in North, East and West Chinese domestic pigs (**Supplemental Fig. S1**). This new view of the odd haplotype pattern on the X-chromosome is well accordant with our previous finding (Ai et al. 2015). Unexpectedly, the distal and proximal region on the Y chromosome of at least 7.7 Mb (hereafter referred to as the SSCY region) displayed two different haplotypes in all tested Euroasian pigs (European pigs, n=30; Chinese pigs, n=71; **Fig. 2A**). The SSCY region was defined as the region from 8.9 to 10.6 Mb and from 39.5 to 43.5 Mb on the VEGA62 version of Y chromosome, meanwhile plus a 2-Mb unmapped contig of Y chromosome. The two haplotypes exhibit a deep divergence as reflected by pairwise nucleotide diversity (**Fig. 2B**) and the phylogenic tree (**Fig. 2C**).

**Figure 2.**
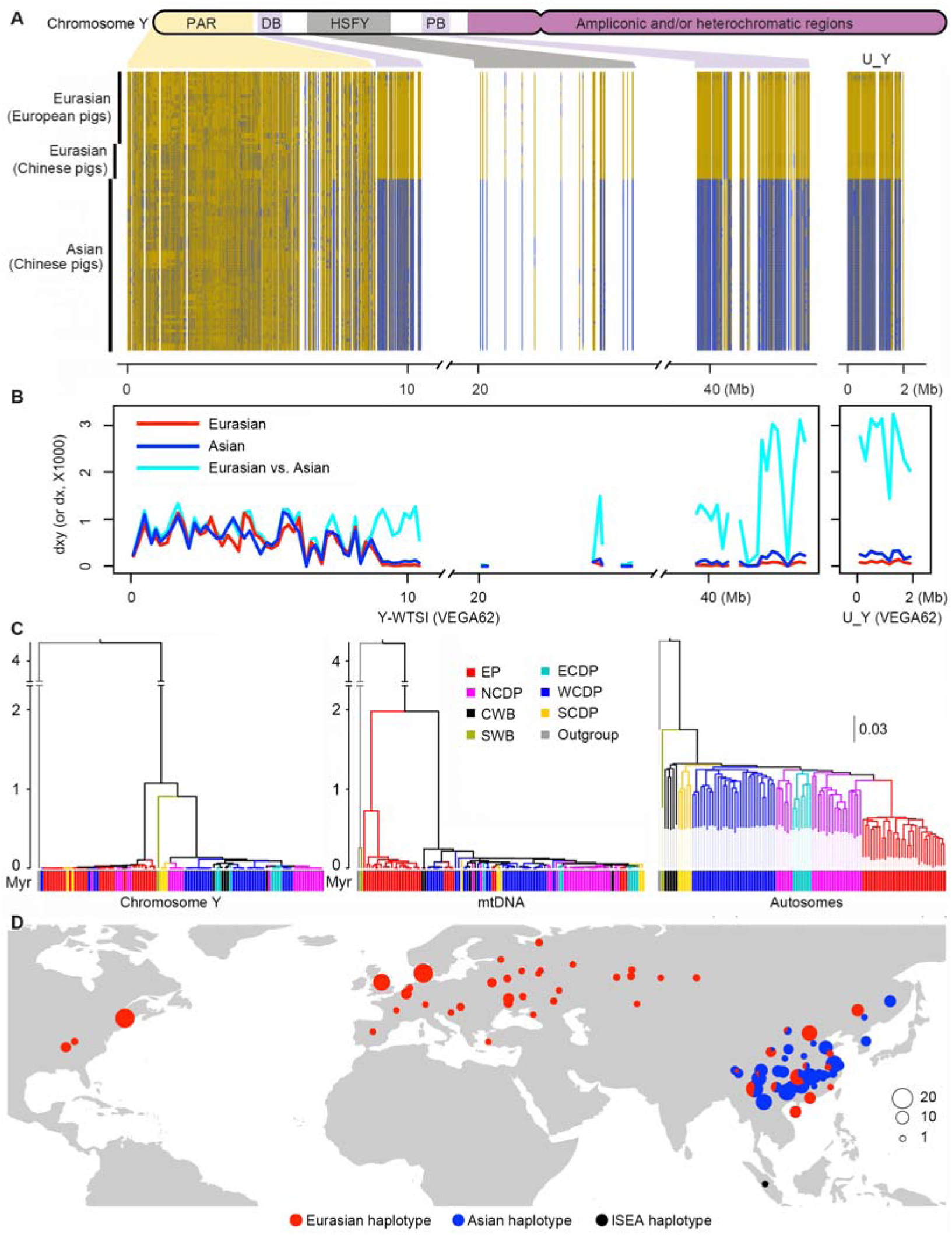
Demographic history of Eurasian pigs based on the data of chromosome Y. (A) The pattern of haplotype on chromosome Y sharing in Eurasian pigs. The haplotypes were reconstructed for each individual using all qualified SNPs on the Y chromosome. Alleles that are identical or different from the ones on the VEGA62 reference genome are indicated by orange or blue, respectively. (B) The plots of *dx* and *dxy* (the number of pairwise differences per site) statistics in a window size of 200 kb on chromosome Y. These statistics were calculated for pigs with the Eurasian or Asian haplotypes. U_Y indicates the unmapped contig on chromosome Y. (C) Bayesian trees of Eurasian pigs constructed using SNP data of the SSCY region and mtDNA, and neighbor-joining tree of these pigs based on autosomal data. Inferred divergence time is shown in Y-axis of the Bayesian trees. The abbreviations of EP, ECDP, NCDP, WCDP, CWB, SCDP and SWB are as in **Figure 1**. *S. verrucosus* (Java warty pig) was set as outgroup. (D) The geographical distribution of the Eurasian (red) and Asian (blue) haplotypes within the proximal and distal region on the Y chromosome (the SSCY region) in Eurasian pigs. The two haplotypes were phased using six tagging SNPs on the porcine 60K Chip (Illumina) within this region. ISEA haplotype (black), the haplotype found in a Sumatran wild boar.

However to our surprise, the distribution pattern of these two haplotypes on the Y chromosome in Eurasian pigs was different from the haplotype pattern on the X chromosome. One haplotype was exclusively found in pigs across China and hence denoted as Asian haplotype; while the other one was found in European pigs and some Chinese pigs from North, South and West China, therefore designated as Eurasian haplotype (**Fig. 2A**). We further investigated the geographical distribution of the two haplotypes in a larger panel of 426 male pigs from 82 diverse populations across Eurasia and America using six tagging SNPs representing the SSCY region (**Fig. 2D** and **Supplemental Table S3**). Again, the Asian haplotype was observed only in East Asian pigs including Chinese pigs, Korean wild boars and Russian Primorsky wild boars; and the Eurasian haplotype was found in European and Chinese pigs, as well as American pigs, which were originated from European pigs (**Fig. 2D**).

### Demography and Evolutionary history inferred by the Y-chromosome

We estimated the divergence time of the haplotypes on the SSCY region using a strict molecular clock implemented in BEAST (Drummond et al. 2012) under the GTR+Γ+I model. In the Bayesian tree, there is a clear split between European pigs with the Eurasian haplotype and Chinese pigs with the Asian haplotype (**Fig. 2C**). Their divergence time was estimated to be 1.07 million years (**Table 1**). However, the time of most recent common ancestors (TMRCA) of European and Chinese pigs with the Eurasian haplotype and of Chinese pigs with Asian haplotype were estimated to be only about 0.10 and 0.14 million years, which are unusually smaller than their splitting time (**Table 1**). And T_MRCA_ of Chinese pigs with the Eurasian haplotype was inferred to be 24k years with 95% confidence intervals from 16k to 30k years (**Table 1**). Also surprisedly, Sumatran wild boar showed much closer relationship to Chinese pigs with Asian haplotype with the splitting time of 0.90 million years (**Table 1**), which is much smaller than the time of 2.1 million years as previously reported using autosomal data (Frantz et al. 2013).

**Table 1.** Time estimates for the TMRCA of phylogenetic nodes of particular interest on the SSCY region.

| Node | Time estimate of<br>$T_{MRCA}$ (Million years) | 95% highest posterior<br>density (HPD) interval |
| --- | --- | --- |
| All the *Sus scrofa* | 1.070 | 0.945-1.156 |
| Sumatran wild boar and Chinese pigs<br>with Asian haplotype | 0.903 | 0.778-0.977 |
| All the pigs with Asian haplotype | 0.136 | 0.111-0.152 |
| All the pigs with Eurasian haplotype | 0.102 | 0.087-0.125 |
| Chinese pigs with Eurasian haplotype | 0.024 | 0.016-0.030 |

To learn more about the evolutionary history of chromosome Y, we further calculated nucleotide variability at the genomic level and within the SSCY region in these sequenced male animals. The SSCY region had a significantly lower level of nucleotide diversity in comparison with autosomes (**Fig. 3**), which is not likely caused by sex-biased genetic drift (**Supplemental Table S4 and S5**) and could be due to the absence of recombination and reduced mutation rates within this region. Moreover, nucleotide variability parameters including segregation sites, *theta* and *Pi* values were all lower in European pigs with the Eurasian haplotype than Chinese pigs with the Asian haplotype (**Fig. 3**, **Supplemental Table S4 and S5**). This could be explained by the fact that European wild boars had suffered a more dramatically decrease of population size than Asian wild boars during the Last Glacial Maximum period (Fig. 1F). Of note, Chinese pigs having the Eurasian haplotype had ∼ six-fold lower values of nucleotide variability than European pigs with this haplotype (**Supplemental Table S4 and S5**), and selection signal was not observed in these Chinese pigs as indicated by Tajima’s D values (**Fig. 3, Supplemental Table S4 and S5**). A reasonable explanation for this observation is that the Eurasian haplotype in Chinese pigs is a European-originated genetic component and has been introgressed into Asian wild boars via an ancient gene flow before domestication approximately 24k years ago. This genetic component contributes to the current gene pool of a proportion of Chinese domestic pigs, possibly via a complicated human-mediated dispersal after pig domestication. Similar to our previous finding of the interspecies introgression on chromosome X (Ai et al. 2015), our ability to detect this ancient male-driven gene flow on chromosome Y is facilitated by the fact that the introgression fragment falls in a region without recombination and low mutation rates and thus can be maintained for a prolonged period. If the introgressed segment had not fallen in such a region, we would likely never have detected the unusual haplotype pattern as recombination and normal mutation rates may quickly degenerate the integrity of introgression fragments. Of note, this evolutionary pattern has not been observed on the Y-chromosome of other mammals like human (Poznik et al. 2016), dog (Shannon et al. 2015) and horse (Wallner et al. 2013).

**Figure 3.**
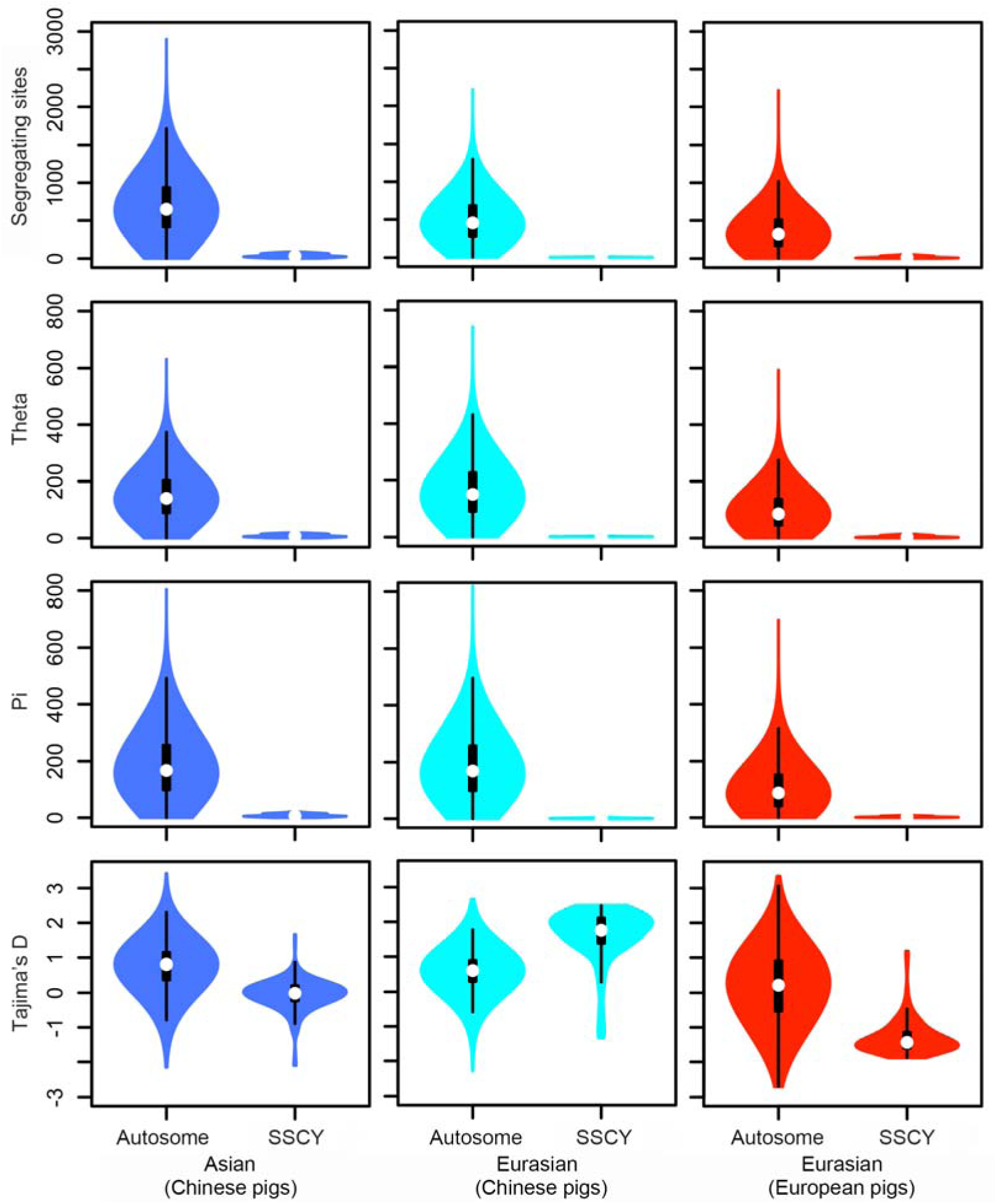
Comparison of nucleotide variability within the proximal and distal region on the Y chromosome and on autosomes. Statistics of segregation sites, theta, Pi and Tajima’s D values were calculated in a window size of 50 kb for European pigs with the Eurasian haplotype, Chinese pigs with the Eurasian haplotype and Chinese pigs with the Asian haplotype on the Y chromosome, respectively.

## Discussion

### Chromosome Y clarifies the evolutionary history of *Sus scrofa*

In the present study, we show a previous unknown evolutionary history of the porcine Y chromosome. In our findings, three key points surprise us. First, only two huge different haplotypes were observed in the distal and proximal blocks of Chromosome Y in a diverse panel of Eurasian pigs (**Fig. 2A**). The genomic and genetic analyses reveal that these regions have two special evolutionary features: no recombination and lower mutation rates, which have not been found in sex chromosomes of other common mammals but only in pigs. Previously we found large region on Chromosome X (48 Mb on the Wuzhishan reference X chromosome; 52 Mb on the Build 10.2 version of Duroc reference X chromosome) harbors the characteristics of low recombination and low mutation rates. We speculated that the enrichment of a 6-kb poly(T) core sequence in the region might contribute to low recombination (Ai et al. 2015). But what exactly contribute to low mutation rates on these regions with large size? Still we don’t know. This unknown biological mechanism, depressing the nucleotide mutation on these regions, merits for further exploration. Second, from the perspective of chromosome Y, we found that the relationship between Sumatran wild boar and pigs from Eurasian continents is much closer than that inferred by autosomes. Previously, Frantz et al. (2013) have observed the discordant phenomenon of conflicting phylogenetic signal between mtDNA and autosomal chromosomes in Sumatran wild boar. They explain this phenomenon by sunda-shelf admixture. Our results of Chromosome Y confirm the hypothesis and make the whole event of inter-specific gene flow more reasonable and much clearer. Third, many Chinese pigs share the same haplotype of chromosome Y with European pigs, which is not consistent with their deep splitting status on autosome and mtDNA. It is well recognized that European and Asian pigs are two geographically and genetically distinct identities (Groenen et al. 2012; Frantz et al. 2013). However, Asian germplasm had been deliberately imported to Europe to improve commercial trait in European breeds during the Industrial Revolution (Bosse et al. 2014), resulting in the fact that ∼30% of the genomes of European commercial pigs are derived from Chinese breeds (Groenen et al. 2012). Here we show another interesting feature of *Sus scrofa* demography, i.e. ancient hybridization between European and Asian wild boars provided an importance source of male genetic components for Asian (Chinese) domestic pigs. These results collectively depict a complex demographic pattern of *Sus scrofa*, significantly advance our knowledge of pig evolutionary history.

### Male-driven gene flow on Chromosome Y is not likely a recent but an ancient event

Given the geographic distance between Europe and Asia, introgression and hybridization between European and Asian domestic pigs were certainly extremely rare before Ferdinand Magellan circumnavigated the globe. More recently, two waves of gene flow between European and Asian domestic pigs have been well documented. One occurred at the onset of Industrial Revolution and early nineteenth centuries. During this period, Asian pigs mainly from South China were introduced into British to improve local breeds, resulting in mosaic modern genomes of British-derived commercial breeds (such as Large White, Landrace and Duroc) that possess ∼30% of Asian haplotypes (White 2011; Groenen et al. 2012). Interestingly, the gene flow orientation shifted afterwards. Since the early twentieth centuries, European improved breeds having desirable performance of lean pork production were imported into China via the Russian, British and German colonies. These breeds including Large White, Berkshire, Duroc and Landrace were introgressed into some Chinese indigenous breeds like Kele and Licha, leaving hybridization signals in the genomes of these breeds (Ai et al. 2014). This raises one possibility that the gene flow observed on chromosome Y could take place after the twentieth centuries via the introgression of European germplasm into Chinese local pigs. This possibility seems to be supported by the observation of very few segregating sites and small Theta and Pi values in Chinese pigs with Eurasian Y haplotype (**Supplemental Table S4**) and of a short branch of these pigs in the Bayesian tree (**Fig. 2C**). However, we can definitely exclude this possibility based on the following four arguments, therefore supporting the conclusion that the male-driven gene flow on chromosome Y is an ancient event before domestication around 24k years ago.

First, nearly all Chinese pig samples used in this study were collected from nucleus herds raised in national conservation farms. These herds are geographically isolated populations. Except for North Chinese domestic pigs, there is a lack of any historical records describing the importation of European breeds into these populations. Indeed, we did not find European ancient lineages in autosomes of most of Chinese domestic pigs in the ADMIXTURE analysis (**Fig. 1C**), which could be detected if there was a recent gene flow between these European and Chinese pigs.

Second, we did not observe any evidence of admixture between European and Chinese domestic pigs on chromosome X. All Chinese pigs with the Eurasian SSCY haplotype do not have the European haplotype of 52 Mb on chromosome X (**Supplemental Fig. S1**). If European germplasm were recently introgressed into Chinese domestic pigs during the last century, it is impossible to retain only Eurasian SSCY segments while completely loss the European chromosome X haplotype in all introgressed individuals via genetic drift or artificial selection within a short time interval of ∼100 years. Unlike European commercial breeds, no intensive selection for specific traits has been documented for Chinese indigenous breeds during the past century.

Third, our mtDNA data show that 20 European pigs (males=9, females=11) including White Duroc, Large White, Landrace and Pietrain have mtDNA of Chinese origin (**Supplemental Table S2**). This is most likely attributable to the recent importation of Chinese breeds to Europe during the 1900s. However, all 134 Chinese pigs (males=71, females=63) lack Europe-originated mtDNA (**Supplemental Table S2**). This also conflicts with the hypothesis of a recent gene flow between European and Chinese domestic pigs.

Forth, we calculated a negative and positive value of Tajima’s D for the SSCY haplotype in European pigs (-1.631) and Chinese pigs (2.350) with the Eurasian haplotype, respectively (**Fig. 3**). For a recently introgressed haplotype, a surplus of rare alleles appearing after the introgression usually creates smaller (i.e. more negative) value of Tajima’s D. Therefore, it is theoretically expect to obtain a more negative value of D in presumably introgressed Chinese pigs, i.e. those with the Eurasian SSCY haplotype. However, the prediction is contradictory to our observation, thus excluding the possibility of a recent gene flow on SSCY.

Together, we conclude that the male-driven gene flow on SSCY is an ancient event, in which hybridization between European and Asian wild boars occurred before domestication (∼24k years ago) and provided an important source of male genetic components for Asian (Chinese) domestic pigs. The very few segregating sites and small values of Theta and Pi in Chinese pigs with the Eurasian SSCY haplotype are most likely due to the unusually evolutionary feature of this region: low mutation rates and no recombination.

## Methods

### Samples and genome sequencing

We sequenced the genomes of 163 Chinese and European pigs, including 24 South Chinese domestic pigs, 33 North Chinese domestic pigs, 36 West Chinese domestic pigs, 33 East Chinese domestic pigs, 6 South Chinese wild boars and 31 European domestic pigs. Of these animals, Chinese pigs were from 16 geographically divergent breeds, European pigs were from 4 commercial breeds (**Supplemental Table S1**), and 59 Chinese pigs have been sequenced in our previous study (Ai et al. 2015).

The genome sequencing was conducted as previously described (Ai et al. 2015). Briefly, genomic DNA was extracted from ear tissues using a standard phenol-chloroform method, and then sheared into fragments of 200-800 bp according to the Illumina DNA sample preparation protocol. These treated fragments were end-repaired, A-tailed, ligated to paired-end adaptors and PCR amplified with 500 bp (or 350 bp) inserts for library construction. Sequencing was performed to generate 100 bp (or 150 bp) paired-end reads on a HiSeq 2000 (or 2500) platform (Illumina) according to the manufacture’s standard protocols.

### SNP calling

We downloaded the genome sequence data of 39 pigs, one African warthog (*Phacochoerus africanus*), one Java warty pig (Sus *verrucosus*) and one Celebes warty pig (*Sus celebensis*) from the NCBI SRA database (https://www.ncbi.nlm.nih.gov/sra). These data were integrated into the sequence data obtained in this study, resulting in a 205-sample high-quality data set (**Supplemental Table S1 and Supplemental Table S2**). Clean reads from all individuals were aligned to the *Sus scrofa* reference genome (build 10.2) using BWA (Li and Durbin 2009). The mapped reads were subsequently processed by sorting, indel realignment, duplicate marking, and low quality filtering using Picard (http://picard.sourceforge.net) and GATK (McKenna et al. 2010). Sequencing coverage and depth of each sample were calculated using genomecov implemented in Bedtools (Quinlan and Hall 2010). A two-round procedure of SNP calling was performed using Platypus (Rimmer et al.2014). In the first round, SNPs were individually called with default parameters, and high-quality called SNPs were merged together. These merged SNPs were treated as known variants to guide the second-round genotyping for all the samples via Platypus with parameters of “‐‐source=KnownVariants.vcf.gz ‐‐minPosterior=0 ‐‐ getVariantsFromBAMs=0”. All SNPs except those on chromosome Y were filtered with the criterion of MAF > 0.001 and SNP call rates > 80%. For SNPs on the Y-chromosome, only male individuals were explored to call SNPs under the criterion of MAF > 0.001 and call rates > 80%.

### Population genetic analysis using autosome data

A total of 47,009,938 qualified SNPs on autosomes were used to calculate genetic distance among all individuals using Plink as previously described (Ai et al. 2013). A neighbor-joining tree was then constructed for all individuals using Neighbor in PHYLIP v3.69 (Felsenstein 2005) and visualized by FigTree software (http://beast.bio.ed.ac.uk/FigTree). Population genetic structure was inferred using the Maximum Likelihood approach implemented in ADMIXTURE v1.20 (Alexander et al. 2009). The ADMIXTURE program was run in an unsupervised manner with a variable number of clusters (K = 2 to 5). Principal component (PC) analysis was conducted using Smartpca in EIGENSOFT v6.0 (Price et al. 2006). To avoid artifacts caused by linkage disequilibrium (LD), we excluded SNPs with r^2^ ≥ 0.2 in the PC analysis.

TreeMix (Pickrell and Pritchard 2012) was employed to infer the patterns of historical splits and mixtures among Eurasian pig populations in the context of *Suidae*, with no migration events and 5000 SNPs grouping together in a LD block. We inferred demographic history of Eurasian pigs using the Multiple Sequentially Markovian Coalescent (MSMC) method (Schiffels and Durbin 2014). To ensure the quality of consensus sequence, we only used representative samples (n=40) of high sequencing depth for each geographic population (n=20) with parameters set as follow: “-p 2*2+50*1+1*4+1*6 ‐‐fixedRecombination” The generation time (g) was set as 5 years, and a standard mutation rate (μ) of 2.5×10^−8^ was used as previously described (Groenen et al. 2012).

### Evolutionary history analysis using chromosome Y data

The bowtie2 software (Langmead and Salzberg 2012) was employed to align filtered clean reads from all male individuals to the chromosome Y reference sequence (VEGA62 version) (Skinner et al. 2016) with the parameter of “‐‐no-mixed ‐‐no-discordant ‐‐no-unal”. Then the two-round SNP calling was performed using Platypus (Rimmer et al. 2014) same as for autosomal SNPs. A total of 49,103 SNPs on chromosome Y passed the criterion of MAF > 0.001 and call rates > 80%. Shapeit2 (Delaneau et al. 2013) was used to phase these SNPs with the parameters of “-X ‐‐burn 14 ‐‐prune 16 ‐‐main 40”. Phased alleles on the distal and proximal blocks were linked together to form sequences, which were then used to reconstruct a phylogenetic tree using BEAST (Tamura et al. 2013) under the GTR+Γ+I model of evolution. Splitting times and 95% highest posterior density intervals in the tree were estimated using a Bayesian Markov chain Monte Carlo method implemented in BEAST (Tamura et al. 2013) with 1,000,000 MCMC samples. The node age of *Sus verucosus* was set to be 4.2 million years (Frantz et al. 2013) as the calibration constraint.

To investigate global distribution of the haplotypes within the distal and proximal region on chromosome Y, we employed six tagging SNPs representing this region from porcine 60K chip in 426 Eurasian pigs from 82 geographically divergent populations. Pairwise nucleotide differences per site within (d_x_) and between (d_xy_) populations were calculated as previously described (Ai et al. 2015). Segregating sites, Theta, Pi and Tajima’s D values were calculated for autosomes and the SSCY region in 50 kb windows at a step size of 25 kb using c++ library of libsequence (Thornton 2003). A coalescence simulation for the SSCY region was performed in Chinese pigs with the Asian haplotype, Chinese pigs with the Eurasian haplotype and European pigs with the Eurasian haplotype using the ms software (Hudson 2002) under demographic model inferred from the MSMC (Schiffels and Durbin 2014) results without recombination.

## Data access

The raw sequence reads from this study have been submitted to the NCBI Sequence Read Archive (SRA; http://www.ncbi.nlm.nih.gov/sra) under accession number SUB2302970.

## Acknowledgements

This study is financially supported by the National Swine Industry and Technology system of China (nycytx-009), Innovative Research Team in University (IRT1136), and the Natural Science Foundation of Jiangxi Province (20152ACB21001).

## Author’s contributions

L.H., J.R. and H.A. designed the study. H.A. and J.R. designed the bioinformatics analysis process. H.A. performed the analyses of bioinformatics, population genetics and evolutionary history. B.Y., J.M., and Z.Z. performed sample collection. W.L. and H.A. performed mapping of raw sequencing reads. H.A., J.R. and L.H. wrote and revised the paper.

## Supplemental materials

### Supplemental Table S1

Samples used in this study. See excel file “Supplemental Table S1.xlsx”.

### Supplemental Table S2

Sequencing statistics of 205 samples used in this study. See excel file “Supplemental Table S2.xlsx”.

### Supplemental Table S3

The geographical distribution of the two haplotypes in a large panel of 426 male pigs from the global. See excel file “Supplemental Table S3.xlsx”.

### Supplemental Table S4

Z-test showing that gene drift is not likely a cause of the two haplotypes on chromosome Y in Eurasian pigs. See excel file “Supplemental Table S4.xlsx”.

### Supplemental Table S5

Coalescent simulations showing that gene drift is not likely a cause of the two haplotypes on chromosome Y in Eurasian pigs. See excel file “Supplemental Table S5.xlsx”.

**Supplemental Fig. S1.**
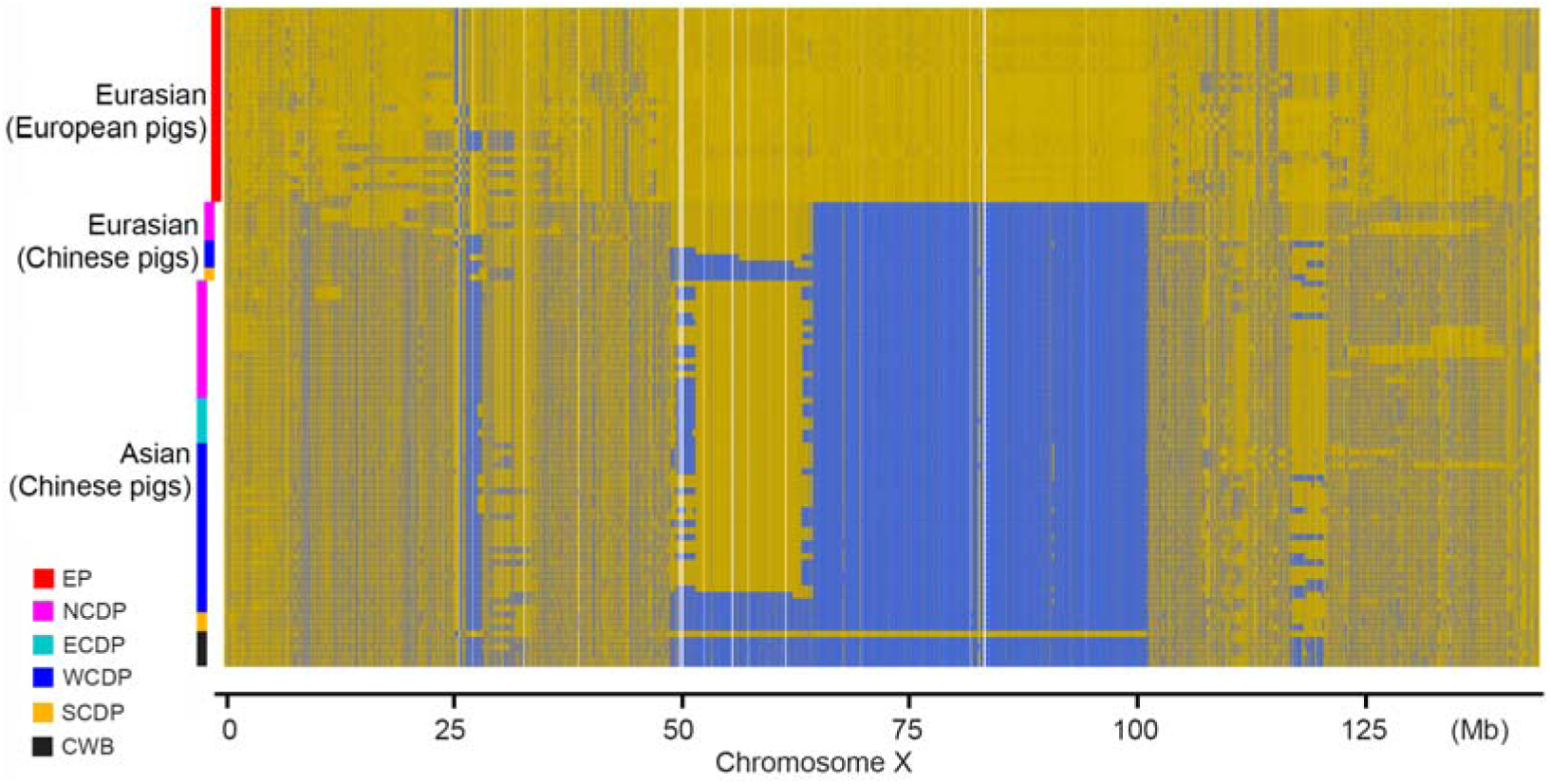
The pattern of haplotype on the X chromosome sharing in Eurasian pigs. The haplotypes were reconstructed for each individual using all qualified SNPs on this chromosome. Alleles that are identical or different from the ones on the Duroc reference genome are indicated by orange or blue, respectively. The abbreviations of EP, NCDP, ECDP, WCDP, SCDP and CWB are as in **Figure 1**.

